# Despite egg-adaptive mutations, the 2012-13 H3N2 influenza vaccine induced comparable antibody titers to the intended strain

**DOI:** 10.1101/158550

**Authors:** Sarah Cobey, Kaela Parkhouse, Benjamin S. Chambers, Hildegund C. Ertl, Kenneth E. Schmader, Rebecca A. Halpin, Xudong Lin, Timothy B. Stockwell, Suman R. Das, Emily Landon, Vera Tesic, Ilan Youngster, Benjamin Pinsky, David E. Wentworth, Scott E. Hensley, Yonatan H. Grad

## Abstract

**Background:** Influenza vaccination aims to prevent infection by influenza virus and reduce associated morbidity and mortality; however, vaccine effectiveness (VE) can be modest, especially for subtype A/H3N2. Failure to achieve consistently high VE has been attributed both to mismatches between the vaccine and circulating influenza strains and to the vaccine's elicitation of protective immunity in only a subset of the population. The low H3N2 VE in 2012-13 was attributed to egg-adaptive mutations that created antigenic mismatch between the intended (A/Victoria/361/2011) and actual vaccine strain (IVR-165).

**Methods:** We investigate the basis of the low VE in 2012-2013 by evaluating whether vaccinated and unvaccinated individuals were infected by different viral strains and assessing the serologic responses to A/Victoria/361/2011 and the IVR-165 vaccine strain in an adult cohort before and after vaccination.

**Results:** We found no significant genetic differences between the strains that infected vaccinated and unvaccinated individuals. Vaccination increased titers to A/Victoria/361/2011 as much as to IVR-165. These results are consistent with the hypothesis that vaccination served merely to boost preexisting cross-reactive immune responses, which provided limited protection against infection with the circulating influenza strains.

**Conclusions:** In contrast to suggestive analyses based on ferret antisera, low H3N2 VE in 2012-13 does not appear to be due to the failure of the egg-adapted strain to induce a response to the intended vaccine strain. Instead, low VE might have been caused by the emergence of antigenically novel influenza strains and low vaccine immunogenicity in a subset of the population.

## Introduction

Vaccination is one of the most important public health interventions to mitigate the annual threat of influenza. Each season, an estimated 4% of the <65-year age group seek outpatient care for influenza-related symptoms, and 12,000-56,000 people die from influenza infection in the US [1, 2]. Vaccination in the US population has been increasing, with estimates from 2016 ranging from 34.3% in the 18-49-year age group to 56.6% in the >65-year age group [3]. However, the protection provided by vaccination continues to be modest, particularly against the H3N2 viruses, with a recent meta-analysis reporting pooled vaccine effectiveness (VE) of 33% (95% CI 26-39) for A/H3N2, 61% (57-65) for A/H1N1pdm09, and 54% (46-61) for B [4].

What explains this modest effectiveness, particularly for the H3N2? A common explanation is that the vaccine and circulating strains are antigenically different, implying that vaccination protects against the vaccine strain but is less protective against antigenically distinct circulating strains. Meta-analyses have indicated that VE to antigenically variant viruses is worse than to matched viruses [4, 5]. Mismatches can arise from inaccurate prediction of the future strains that will dominate a season and the significant antigenic diversity of co-circulating strains [6–8] as well as mutations associated with egg adaptation that are acquired during, and facilitate, vaccine production by most large-scale manufacturers [9].

Antigenic mismatch between the vaccine and circulating strains may have lowered VE during the H3N2-dominated 2012-13 season, when influenza VE was estimated to be 39% (95% CI, 29-47) [10]. The limited effectiveness was attributed to mutations in the intended H3N2 vaccine strain, A/Victoria/361/2011 (Vic/361), in several epitopes that arose during growth in eggs, yielding an egg-adapted vaccine strain used by vaccine manufacturers (IVR-165) [11]. Although some of these mutations occurred in the head of the hemagglutinin (HA) near the receptor-binding site (H156Q, G186V, S219Y) and may thus be especially immunogenic [12], it is notable that the estimated VE for the season was typical for recent A/H3N2 seasons.

Low VE might also have arisen from especially poor vaccine-induced protection in particular host subpopulations [13, 14]. Differences in VE by age [4, 10] have led to speculation that age-associated differences in infection and vaccination history [15, 16] affect protection [17, 18]. Notably, naive ferrets infected with IVR-165 developed titers that were 16-to 32-fold lower against Vic/361 compared to IVR-165, but naive ferrets infected with Vic/361 developed similar titers to both strains [11]. People previously uninfected with Vic/361-like viruses who were immunized with the IVR-165 vaccine strain might have similarly distinct responses, poorly cross-reacting with Vic/361 and perhaps reacting differently with circulating clades compared to people with immunity to Vic/361. It is unclear, however, if a poorly cross-reacting response would have been induced if the vaccine recipients had already been exposed to Vic/361-like viruses. Previous exposures to other strains might expand the range of possible responses to a vaccine.

To investigate the basis of the low VE against H3N2 in 2012-2013, we first examined whether vaccinated and unvaccinated individuals were infected by different viral strains. If vaccinated people were infected with antigenically distinct strains, then low VE may be partially attributable to mismatch of some kind. We then evaluated the serologic responses to Vic/361 and IVR-165 in an adult cohort before and after vaccination. If low VE arose from mismatch due to egg adaptations in IVR-165, the expectation is that we should see a robust response to IVR-165 but not to Vic/361.

## Methods

### Dataset

The dataset is comprised of influenza A/H3N2 samples from 2012-13 collected for this study and sequences collated from online databases. The samples collected for this study were from clinical specimens identified as positive for influenza in the microbiology labs in three hospitals in Boston, MA (Brigham and Women’s Hospital; Massachusetts General Hospital; Boston Children’s Hospital), one in Chicago, IL (University of Chicago Hospital), and one in Santa Clara, CA (Stanford University Hospital). The CT value and the type and subtype of influenza were recorded, when available. Patient demographics (including gender and age) and vaccination status were documented based on review of the medical record. The study was performed with IRB approval from each of the participating institutions (Partners Healthcare IRB protocol 2013-P-000097/1; Boston Children’s Hospital IRB protocol IRB-2013-P00007299); Stanford IRB protocol 27044; University of Chicago IRB protocol 13-0149). Sequences were also obtained from the NCBI influenza Virus Database (https://www.ncbi.nlm.nih.gov/genomes/FLU/Database/nph-select.cgi?go=database). The query specified full-length HA H3N2 specimens from the USA from 1 September 2012 to 1 June 2013. To these sequences, we added 23 partial HA sequences from Dinis et al. [19] for which the vaccination status of the infected individual was reported. We annotated the genome sequences by date and location of isolation, when available, as well as the vaccination status of the infected individuals [10, 19].

### Sequencing of the influenza genomes

The complete genomes of the influenza A viruses collected were sequenced as part of the NIH/NIAID sponsored Influenza Genome Sequencing Project. Viral RNA was isolated using the ZR 96 Viral RNA kit (Zymo Research Corporation, Irvine, CA, USA). The influenza A genomic RNA segments were simultaneously amplified from 3 μL of purified RNA using a multi-segment RT-PCR strategy (M-RTPCR) [20, 21]. The M-RTPCR amplicons were used as templates for Nextera Library construction and libraries were sequenced using the MiSeq v2 platform (Illumina, Inc., San Diego, CA, USA). The sequence reads from the MiSeq were sorted by barcode, trimmed, and non-influenza sequences were removed. The NGS reads were then mapped to the best matching reference virus using the clc_ref_assemble_long program. At loci where NGS platforms agreed on a variation (as compared to the reference sequence), the reference sequence was updated to reflect the difference. A final mapping of all next-generation sequences to the updated reference sequences was then performed.

### Phylogenetic analysis

HA sequences from the full set of H3N2 influenza strains from 2012-13 were aligned using MUSCLE [22], and a maximum likelihood tree generated using RAxML [23] with default options. Phylogenies and metadata were visualized in Phandango (http://jameshadfield.github.io/phandango/). To determine the association of vaccination status and phylogeny, the Fitz and Purvis *D* statistic was calculated in R using phylo.d from the caper package [24], using a maximum likelihood tree comprised of the isolates with known vaccination status.

### Human subjects and sera

As part of a cohort study [25–27], blood was collected from adult subjects from the Durham-Raleigh-Chapel Hill, NC area in the Duke Clinical Research Unit, Duke University Medical Center Durham, NC.

### Virus propagation and characterization

We used reverse-genetics to produce reassortant viruses expressing either the Vic/361 or IVR-165 HAs. These viruses possessed the same Vic/361 NA and 6 internal protein coding genes from the A/Puerto Rico/8/1934 virus. The HAs of Vic/361 and IVR-165 differed at residues 156 (H156Q), 186 (G186V), and 219 (S219Y). We propagated viruses for 2 days using MDCK-SIAT1 cells and we used standard Sanger sequencing to verify that other HA mutations did not predominate after virus propagation. Contemporary H3N2 viruses often acquire NA mutations during *in vitro* propagation, and these mutations can confound HAI assays [28]. To verify that agglutination of guinea pig erythrocytes of our viral preps was HA-mediated, we completed agglutination assays in the presence or absence of 10uM oseltamivir, a compound that binds in the sialic acid binding site of NA.

### Hemagglutination inhibition (HAI) assays

Sera samples were pre-treated with receptor-destroying enzyme (Key Scientific Products Inc) and HAI titrations were performed in 96 well round bottom plates (BD). Sera were serially diluted twofold and added to 4 agglutinating doses of virus in a total volume of 100 ul. Guinea pig erythrocyte solution (12.5μl; 2% vol/vol) (Lampire) were added to sera/virus mixtures. Agglutination was read out after incubating erythrocytes for 60 min at room temperature. HAI titers were expressed as the inverse of the highest dilution that inhibited 4 agglutinating doses of guinea pig erythrocytes.

## Results

### Vaccine status did not affect infection risk with different clades of H3N2

To assess whether vaccinated and unvaccinated individuals were infected by genetically related viruses, we collated 423 influenza A/H3N2 sequences from individuals with known vaccination status infected during the 2012-13 season in the US. This dataset comprised 316 specimens sequenced for the purposes of this study and two published collections [10, 19] (Supplemental Table S1). Analysis of the HA sequences revealed no phylogenetic clustering by vaccination status (Figure 1), with the Fritz and Purvis [29] estimated *D* statistic of 0.93, consistent with a random association between vaccination status and the phylogeny. This is in keeping with a previous observation using a smaller dataset [19].

**Figure 1.**
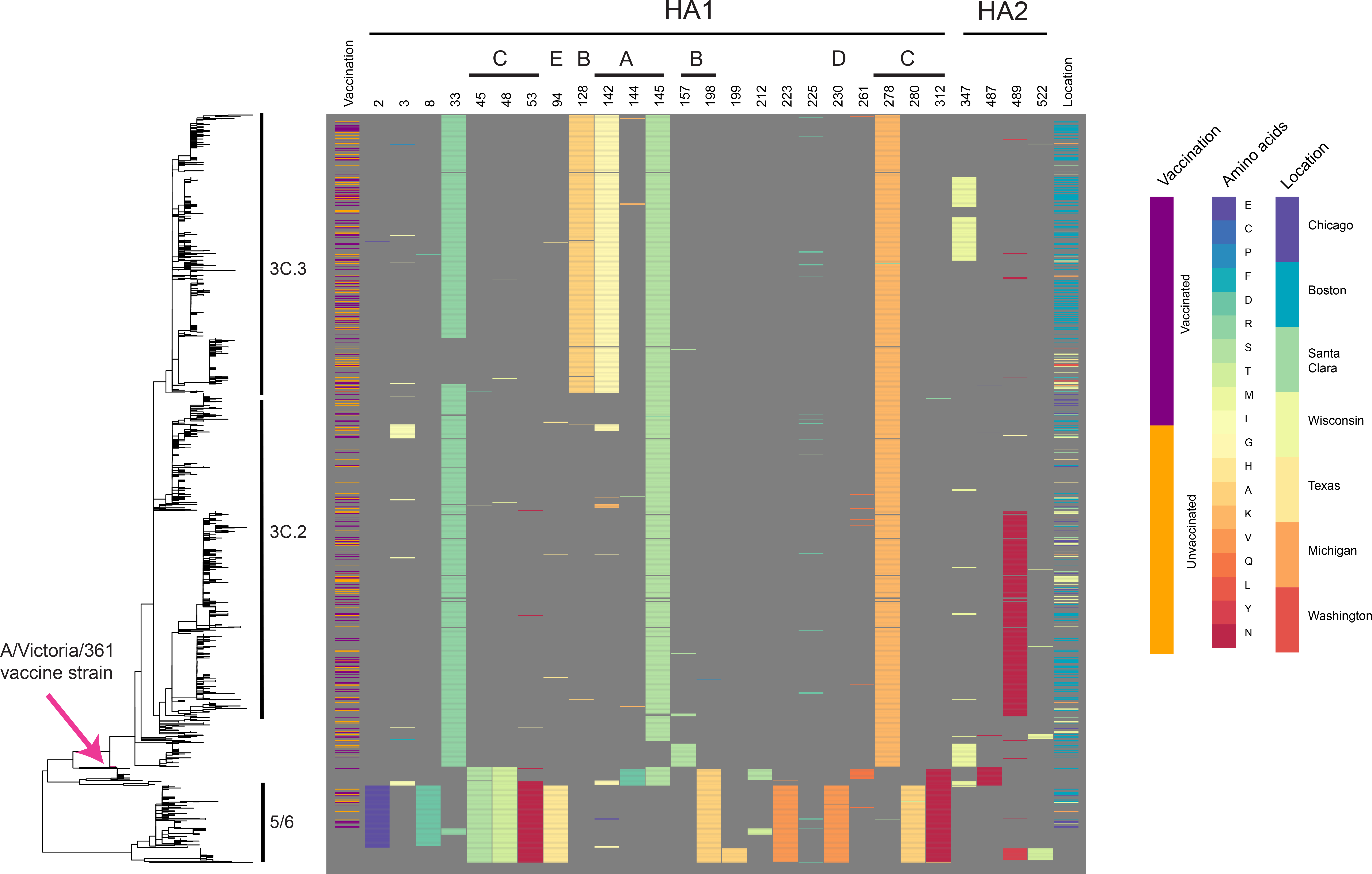
Maximum likelihood phylogeny of HA sequences from 1339 influenza A/H3N2 isolates collected from North America during the 2012-13 influenza season. Vaccination status of the individuals from whom the isolates were collected is noted (purple=vaccinated; orange=unvaccinated; blank=unknown vaccination status). Amino acid sites in which 20 or more of the 1339 specimens differed from the vaccine strain IVR-165 are noted, with the amino acids colored according to the key, and annotated according to their location in HA1, HA2, and predicted epitope sites (A-E). The location from which the isolates were collected is color coded according to the key.

Meaningful differences between the strains infecting vaccinated and unvaccinated individuals may be focused in sites within HA that impact antigenicity and avidity [12, 30] rather than the whole HA sequence. We assembled a dataset of 1339 complete HA sequences from the USA from 2012-13 (Supplemental Table S2), including the same set of 423 sequences described above, and determined the amino acid sites in which 20 or more specimens differed from the vaccine strain IVR-165, yielding 23 sites in HA1 and 4 sites in HA2 (HA2 included for completeness; Figure 1). Based on these 27 sites, we identified a total of 33 haplotypes present in the isolates from individuals with known vaccination status, with frequencies ranging from 1 (n=17) to 122 (n=1) (Supplemental Table S3). For the three most abundant haplotypes, we evaluated whether the samples from vaccinated individuals were overrepresented. A haplotype containing HA2 V18M trended towards significance (Bonferroni corrected *p* value of 0.1). However, 55 out of 59 of the isolates with this haplotype are from Boston, which has a high ratio of vaccinated to unvaccinated individuals (1.7:1) compared to other sites (0.8:1), indicating even less significance. Notably, this mutation is at the root of a section of clade 3C.3, including 3C.3b, and increased in prevalence in the 2014-15 influenza season (https://www.crick.ac.uk/media/221813/nimr-report-feb2015-web.pdf). The haplotype analysis reveals evidence of convergent evolution, with identical amino acid variants appearing in multiple clades (e.g., N225D appears in clades 3C.3, 3C.2, as well as 5/6). Together, these data suggest that there are no statistically significant differences in haplotypes between vaccination groups in a season that showed the emergence of new HA lineages (3C.2 and 3C.3). Rather, both vaccinated and unvaccinated individuals were infected with antigenically diverged clades.

### Vaccination boosts serum antibody responses to vaccine and wild-type strains equally

Because vaccinated people were infected with similar strains as unvaccinated people, antibody responses induced by the vaccine might have been very weak, or they might have recognized epitopes present in the vaccine strain but absent in circulating strains. It has been proposed that egg-adaptations in the IVR-165 vaccine strain contributed to low VE during the 2012-2013 season [11]. Consistent with previous reports [11] we found that naïve ferrets infected with IVR-165 generated antibodies that reacted poorly with the wild-type Vic/361/11 strain (Supplemental Table S4). However, we found that vaccinated adult humans produced antibodies that recognized both Vic/361/11 and IVR-165 (Fig 2 and Supplemental Table S5). For these experiments, we completed HAI assays using sera from 62 adults before and after immunization with IVR-165. Nearly half of subjects had pre-vaccination titers ≥40 to Vic/361 or IVR-165 (Figure 2a,c). After vaccination with IVR-165, titers to the vaccine strain more than doubled (fold-change mean=2.54, SD=3.16) but increased ≥4-fold in only 8% of subjects. Strikingly, despite variation in individuals’ responses to IVR-165, there was a strong linear correlation between subjects’ antibody increases to IVR-165 and Vic/361 (Pearson’s correlation = 0.88 [CI 0.81, 0.93], *p* < 10^−15^) (Figure 2b). Regressing the response to Vic/361 against that to IVR-165 yielded a slope of 0.86 (0.056 SE). Thus, regardless of strength, responses to the vaccine strain induced almost the same magnitude change in response to the wild-type strain, suggesting that the vaccine induced antibodies that cross-reacted with the wild-type strain, and effectively *only* antibodies that cross-reacted with the wild-type strain. The strong correlation between pre-vaccine titers to Vic/361 and IVR-165 (Pearson’s correlation = 0.92 [CI 0.88, 0.95], Fig. 2d) suggests that these antibodies predated exposure to IVR-165. Almost none of the adults responded like naive ferrets, which developed four-fold greater responses to IVR-165 than Vic/361 after vaccination with IVR-165: only one human subject had fold-changes to IVR-165 that were more than two-fold greater than changes to Vic/361.

**Figure 2.**
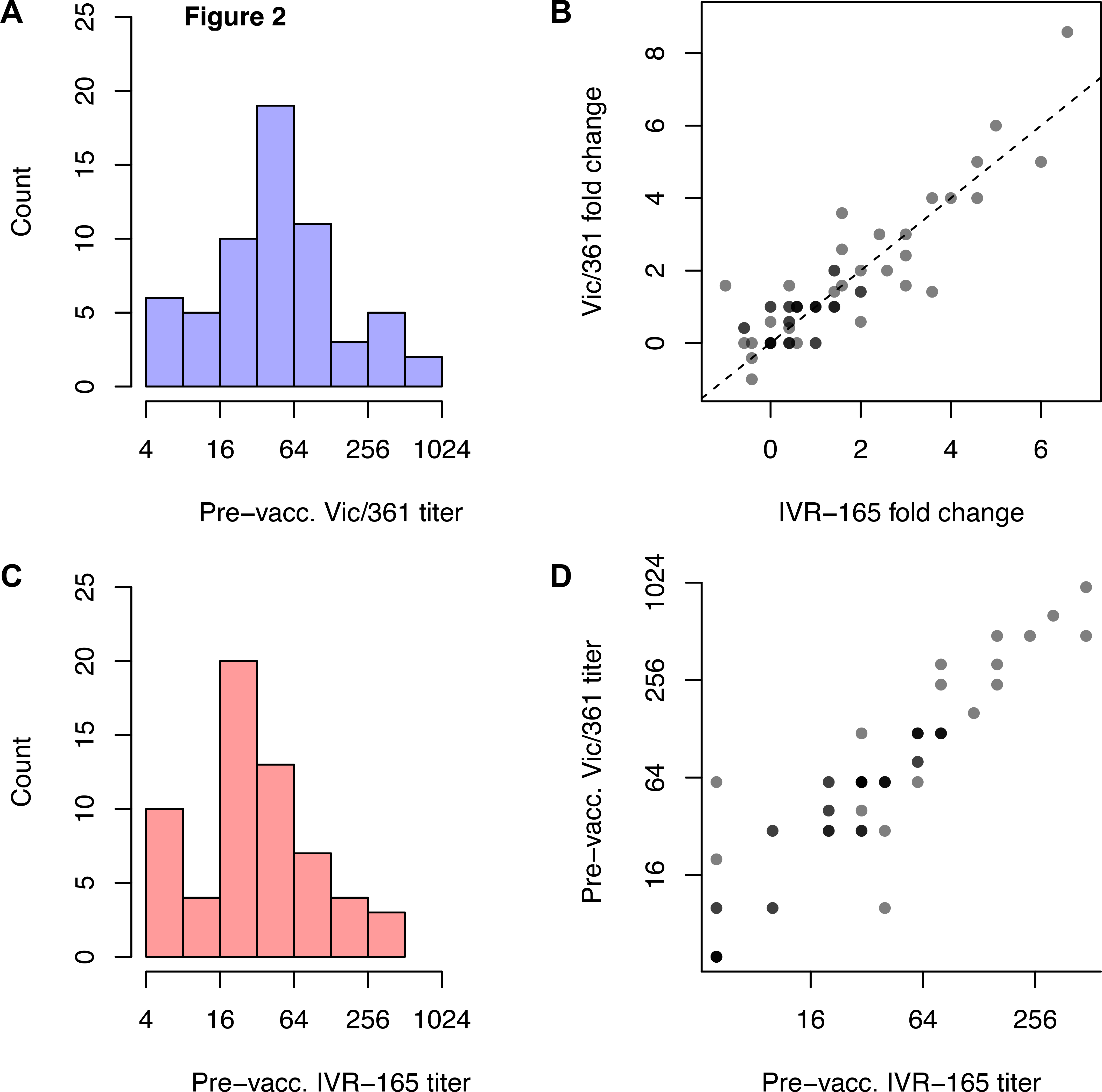
(A) Pre-vaccination log titers to the wild-type Vic/361 (WT) and (C) egg-adapted (IVR-165) strains. (B) Fold changes in responses to each strain after vaccination. Points are semitranslucent; darker points represent multiple individuals. (D) Correlation between prevaccination titers to Vic/361 and IVR-165.

## Discussion

Influenza VE measures how well the vaccine protects against diverse influenza viruses that circulate over a season. Given the high morbidity of H3N2 infections [31, 32], there is an urgent need to understand this subtype’s modest VE. The two general explanations for why the vaccine fails to achieve high effectiveness are (1) poor vaccine match, where “match” refers to the antigenic similarity between the vaccine and circulating influenza strains [4, 5], and (2) heterogeneous responses to influenza vaccination, such that the vaccine elicits protective immunity in only a subset of the population [13, 14].

For the 2012-13 influenza season, the H3N2 VE was estimated to be 39% [10]. A proposed explanation for the low VE focused on three epitope-site mutations that arose during egg-adaptation, suggesting that these mutations resulted in antigenic mismatch between the vaccine and circulating influenza strains [11]. Antigenic data from ferrets support the contention that vaccination with the egg-adapted variants such as IVR-165 yield antibodies that react poorly with the intended Vic/361 vaccine strain. Under this model, the observed VE comes from mismatch with Vic/361, which presumably correlates with the degree of mismatch with circulating strains. In theory, the VE would have been higher without the egg-adaptation mutations, although some mismatch between Vic/361 and circulating strains might have led to imperfect effectiveness.

In contrast with the ferret data, we found that human responses to Vic/361 and IVR-165 strains before and after vaccination with IVR-165 were similar and highly correlated. Human exposure to the vaccine containing IVR-165 induced comparable responses to Vic/361. The differences between the specificity of antibodies elicited in ferrets and humans in our study is likely due to prior H3N2 exposures in humans. There is extensive evidence that B cell responses to influenza strains often evolve from preexisting responses [16, 17, 33–36].

We also observed heterogeneity in the extent of response to vaccination, with some individuals having persistently low HAI titers to the vaccine strain despite vaccination. There is accumulating evidence that increased influenza exposure and older age, which are partly confounded variables [37], are associated with lower rises in titer after vaccination. However, it is unclear why some individuals have lower titers than others after seemingly similar exposure histories, an important problem because titers correlate with protection.

There were no significant differences at the haplotype level between the strains that infected vaccinated versus unvaccinated people. Although this test lacks power, since vaccine recipients who became infected might have been those with a poor response to the vaccine, the results are consistent with the hypothesis that vaccination served merely to boost preexisting Vic/361-like responses, which failed to protect against emerging co-circulating clades.

Together, these data argue that the low H3N2 VE in the 2012-13 season was not primarily attributable to egg-adaptation mutations in the H3N2 vaccine strain. Other explanations, including a general mismatch between Vic/361 and circulating strains, as well as widely variable, and still inexplicable, immunological responses to the vaccine in the vaccinated population, might thus explain the low VE.

## Acknowledgements

The authors also thank Nadia Fedorova and Susmita Shrivastava for technical expertise in support of the viral assembly and submission pipeline for this project. The data for this manuscript was generated while DEW was employed at JCVI. The opinions expressed in this article are the authors own and do not reflect the views of the Centers for Disease Control, the Department of Health and Human Services, or the United States government.

## Data availability

All sequences are available in GenBank; accession numbers provided in Supplementary Material.

